# Fast quantification of uncertainty in non-linear diffusion MRI models for artifact detection and more power in group studies

**DOI:** 10.1101/651547

**Authors:** R.L. Harms, F.J. Fritz, S. Schoenmakers, A. Roebroeck

**Affiliations:** Dept. of Cognitive Neuroscience, Faculty of Psychology & Neuroscience, Maastricht University, the Netherlands

**Keywords:** Uncertainty estimates, Variances, Diffusion MRI, Microstructure, Fisher Information Matrix (FIM), Cramér Rao Lower Bound (CRLB)

## Abstract

Diffusion MRI (dMRI) allows for non-invasive investigation of brain tissue microstructure. By fitting a model to the dMRI signal, various quantitative measures can be derived from the data, such as fractional anisotropy, neurite density and axonal radii maps. The uncertainty in these dMRI measures is often ignored, while previous work in functional MRI has shown that incorporating uncertainty estimates can lead to group statistics with a higher statistical power. We propose the Fisher Information Matrix (FIM) as a generally applicable method for quantifying the parameter uncertainties in non-linear diffusion MRI models. In direct comparison with Markov Chain Monte Carlo sampling, the FIM produces similar uncertainty estimates at lower computational cost. Using acquired and simulated data, we then list several characteristics that influence the parameter variances, like data complexity and signal-to-noise ratio. In individual subjects, the parameter standard deviations can help in detecting white matter artifacts as patches of relatively large standard deviations. In group statistics, we recommend using the parameter standard deviations by means of variance weighted averaging. Doing so can reduce the overall variance in group statistics and reduce the effect of data artifacts without discarding data from the analysis. Both these effects can lead to a higher statistical power in group studies.

## 1 Introduction

Diffusion Magnetic Resonance Imaging (dMRI) allows for non-invasive investigation of brain tissue microstructure. By fitting a dMRI model to each voxel, various quantitative measures can be derived from the data, such as fractional anisotropy (Basser et al., 1994), neurite density (Zhang et al., 2012) and axonal radii maps (Assaf & Pasternak, 2008; Alexander et al., 2010). These quantitative measures can be used in statistical group analysis. For example, tract-based spatial statistics (TBSS) is a popular approach to group analysis of fractional anisotropy measures (Smith et al., 2006). More often than not, these approaches (including TBSS) ignore the uncertainty in the quantitative measures. In functional magnetic resonance imaging, previous work has shown that incorporating uncertainty estimates can lead to group statistics with a higher statistical power (Chen et al., 2012; Woolrich et al., 2004). For linear diffusion models, a method for computing and using uncertainty estimates has been shown before (Sjölund et al., 2018), but this has not yet been generalized to non-linear diffusion models like NODDI (Zhang et al., 2012) and CHARMED (Assaf & Basser, 2005).

Previous work in quantifying the parameter uncertainties include Markov Chain Monte Carlo (MCMC) (Behrens et al., 2003; Wegmann et al., 2017; Gu et al., 2017) and bootstrapping (Jones, 2003; Chung et al., 2006; Whitcher et al., 2008) methods. Of these two techniques, bootstrapping is often not applicable as it is either model specific (Whitcher et al., 2008) or requires very specific additional MRI measurements (Jones, 2003; Chung et al., 2006) which are often not acquired in diffusion MRI datasets. MCMC on the other hand can readily be extended to all microstructure models, but often requires long computation times, even with parallel processing on graphical processing units (Harms & Roebroeck, 2018).

We propose the Fisher Information Matrix (FIM) as a generally applicable method for quantifying the parameter uncertainties in non-linear diffusion MRI models. The FIM allows for estimating the local variances around the maximum likelihood point estimate, which is the point estimate typically used in group statistics. Computing the FIM is a relatively fast operation, requiring only a few additional model evaluations. In other fields, like for example astrophysics, the Fisher Information Matrix is already recognized as a useful tool for quantifying the uncertainty in parameter estimates (Vallisneri, 2008; Rodriguez et al., 2013). In diffusion MRI, the FIM has been applied before, but only specific to the multi-Tensor model (Versteeg et al., 2018) and has not yet been generalized to all non-linear microstructure models.

The Fisher Information Matrix can additionally be used to compute the Cramér Rao Lower Bound (CRLB; Rao, 1945; Cramer, 1946), if and only if the true parameters are known (Kay, 1993). For example, in simulation studies the CRLB can function as a ground truth lower bound on the estimable variances, thereby indirectly evaluating the performance of the maximum likelihood routines (Kay, 1993). Although in brain data the FIM can be interpreted as an approximation to the CRLB, we follow the results in astrophysics and only interpret the FIM as a measure of uncertainty around the estimated parameters (Vallisneri, 2008).

We first compare the uncertainty estimates from the Fisher Information Matrix to those of MCMC, using multiple datasets and multiple dMRI microstructure models. We then investigate several data and model characteristics that can influence the parameter variances, like data complexity and Signal-to-Noise Ratio (SNR). In the end, we discuss the use of uncertainty estimates in white matter artifact detection (e.g. detecting fat saturation) and show how weighted averaging could lead to an increase in power in group studies.

## 2 Methods

### 2.1 Parameter distribution estimates

We compare two different methods for summarizing the parameter posterior distributions of a single voxel, a frequentist method using Maximum Likelihood Estimation (MLE) and the Fisher Information Matrix (FIM) and a Bayesian method using Markov Chain Monte Carlo (MCMC) (see figure 1 for a schematic overview). With both methods we summarize the voxel-wise posteriors as a point estimate with a corresponding standard deviation (std.).

**Figure 1:**
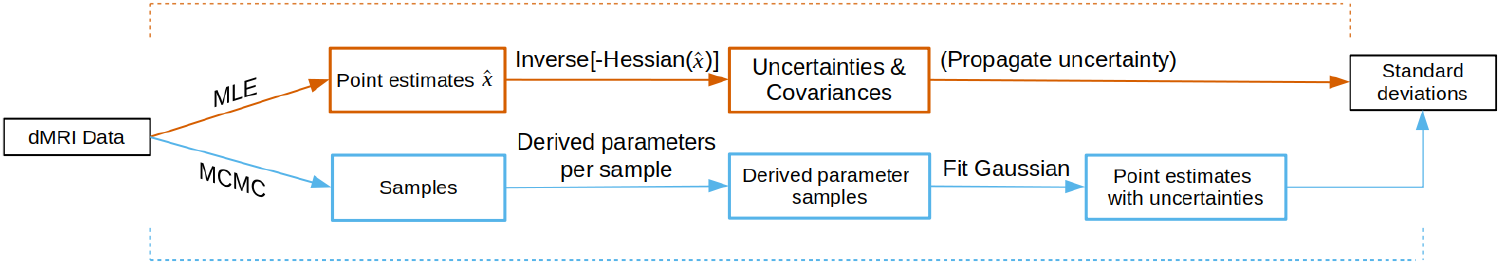
The uncertainty computation methods for both the Maximum Likelihood Estimation (MLE) and Markov Chain Monte Carlo (MCMC) methods.

In the first method we use the Powell optimization routine (Powell, 1964; Harms et al., 2017) to get an MLE parameter point estimates. We estimate the standard deviations around those point estimates using the theory of the FIM. Standard deviations in derived parameter maps (e.g. Tensor Fractional Anisotropy) can be obtained by propagating the uncertainty of the model parameters. We refer to this method as MLE+FIM.

The second methodology uses MCMC sampling to approximate the full posterior distribution, using the Adaptive Metropolis Within Gibbs routine as discussed in (Harms & Roebroeck, 2018). From these samples we summarize the posterior distribution using a mean and standard deviation, as done before in before in dMRI modeling (Behrens et al., 2003; Sotiropoulos et al., 2013; Wegmann et al., 2017). Uncertainties in derived parameter maps can be obtained by computing the derived parameter maps at every sampled point and summarizing the result. We refer to this method as MCMC.

The MLE+FIM provides a local variance around a mode while MCMC provides a global variance around the mean. As such, these methods are only comparable if the posterior is unimodally Gaussian distributed, since then the mean equals the mode. As in previous work (Behrens et al., 2003; Sotiropoulos et al., 2013; Wegmann et al., 2017), we assume the posteriors to be unimodally Gaussian distributed.

This assumption may not necessarily hold. For example, multi-modality could arise when fitting a single fiber model to a crossing fiber voxel. In such cases, different post-processing would be required on the MCMC samples to correctly reflect the parameter variances. The FIM would be less sensitive to this issue since the FIM provides only local variances estimates. That is, the MLE would choose one mode of the distribution and the FIM would provide a local variance estimate around the chosen mode. This issue could also be circumvented by applying appropriate model selection to every voxel.

Non-Gaussian distributions can happen near parameter boundaries. For instance, very low (close to zero) or very high (close to one) compartment volume fractions can lead to a truncated posterior. In such cases the FIM no longer applies. For MCMC different post-processing would be required, like fitting a truncated normal distribution to the posterior. This could again be solved by appropriate model selection. We take no special precautions for these boundary effects and assume these to not be present in white matter.

Nevertheless, we expect most posteriors to be unimodally Gaussian distributed. This assumption is also supported by two theoretical arguments. First, if the model is suitable to describe the data (e.g. if model selection was successfully applied), the posterior asymptotically approaches a Gaussian distribution (Gelman et al., 2013). Second, according to the central limit theorem, each parameter’s marginal distribution will asymptotically tend to a Gaussian as the number of model parameters increases (Gelman et al., 2013).

#### 2.1.1 Fisher Information Matrix

The observed Fisher Information Matrix is defined as the negative Hessian of the log-likelihood function when evaluated at the maximum likelihood estimate (Pawitan, 2013; Gelman et al., 2013). The inverse of the FIM is an asymptotic estimator of the covariance matrix (Pawitan, 2013; Gelman et al., 2013). Formally, let *l*(**x**) be a log-likelihood function with maximum likelihood estimate 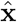. A second order Taylor approximation of *l*(**x**) centered at 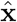 is then given by:

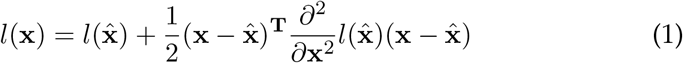

ignoring the higher terms and having dropped the linear term since the first derivative of a function is zero at the mode. Considering the first term, 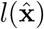, a constant and the second term, 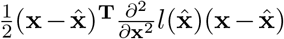, proportional to the logarithm of a normal density, we get the approximation:

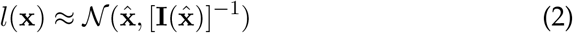

where 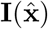 is the observed Fisher Information Matrix:

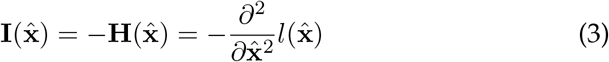

For the Hessian to be positive definite, this theory requires 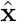 to lie within the boundaries of the parameter space (Gelman et al., 2013). We compute the Hessian numerically (see Appendix A) and its inverse using a direct inverse where possible with a fallback on the (Moore-Penrose) pseudoinverse for ill-conditioned Hessians. Ill-conditioned Hessian can for example arise with parameter estimates lying at a predefined parameter boundary (Gelman et al., 2013).

#### 2.1.2 Uncertainty propagation

Given a function **y** = *f* (***θ***) where *f* (·) is a known function, uncertainty propagation provides the probability distribution of **y** given the probability distribution of ***θ***. For example, we can use this to estimate the standard deviation of a Tensor Fractional Anisotropy (FA) estimate, by propagating the standard deviation estimates of the Tensor diffusivities. We use a first order Taylor expansion linear approximation (Arras, 1998), which states that if ***θ*** is normally distributed with mean ***μ***_***θ***_ and covariance matrix Σ_***θ***_, the distribution of **y** can be approximated as:

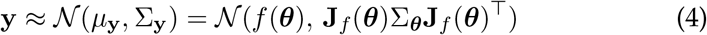

with **J**_*f*_ the Jacobian matrix of *f*. More succinctly, the covariance matrix of **y** = *f* (***θ***) is given by:

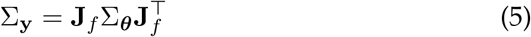

which holds as a generally applicable formula for linear propagation of co-variances (Arras, 1998). In the case of an univariate output *y* = *f* (***θ***), the Jacobian can be formulated as a gradient vector ∇_*f*_, leading to the following expression for the variance in *y*:

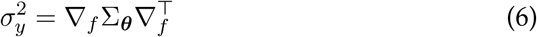

This error propagation technique uses both the variances and the co-variances of all the propagated parameters. Additionally, this technique takes into account the functional form of the propagated function, i.e. if the function is linear or non-linear. The Jacobian or gradient can be computed numerically using finite-differences or can be evaluated at an analytical derivative. We use analytical expressions for all uncertainty propagations. See Appendix B for worked out error propagation examples of the Tensor FA and Ball&Stick Fraction of Stick parameters.

### 2.2 Variance weighted average

Variance weighted averaging makes it possible to include the variances of the data points when computing a mean and standard deviation. For example, the voxel-wise variances discussed earlier can be used in averages of white matter regions within a subject, or in voxel-wise averages over multiple subjects. First, given *n* data points *z*_*i*_, we define the regular mean as:

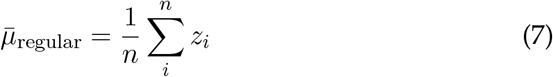

and regular standard deviation as:

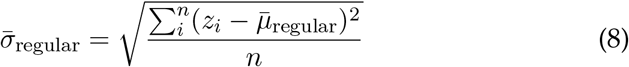

If each data point *z*_*i*_ has a corresponding weight *w*_*i*_, we can compute a weighted mean as:

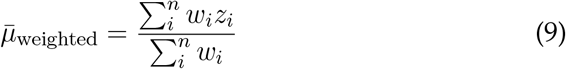

and a weighted standard deviation as:

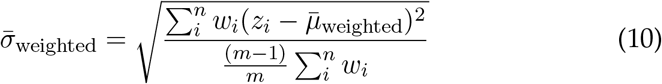

with *m* for the number of non-zero weights, included here to allow for non-normalized weights. It has been shown that the weights that minimize the variance of the weighted average are the reciprocals of the variances of each of the data points *z*_*i*_ (Shahar, 2017). That is, given the variances 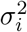 for each *z*_*i*_, the weights that minimize Var(Σ_*i*_ *w*_*i*_*z*_*i*_) is given by:

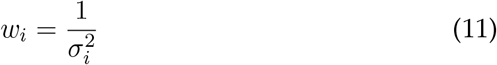

Incidentally, these weights are also the maximum likelihood estimator of the weighted mean and variance under the assumption that the data points *z*_*i*_ are independent and normally distributed with the same mean (Cochran, 1937).

### 2.3 Diffusion microstructure models

To capture the variety of microstructure models in diffusion MRI we chose four different models, the Tensor (Basser et al., 1994), Ball&Stick (Behrens et al., 2003), Bingham-NODDI (Tariq et al., 2016) and CHARMED (Assaf et al., 2004) models. The Tensor model is the oldest diffusion MRI model and still sees widespread usage in the literature. From the Tensor we derive the Fractional Anisotropy (FA) quantity. The Ball&Stick model (Behrens et al., 2003) is the first multi-compartment model and is often used as local estimator for tractography. To delineate multiple fiber orientations, the Ball&Stick model can feature multiple Stick compartments, but always with a single Ball compartment. To differentiate between the Ball&Stick models with one or more Stick compartments, we denote the specific Ball&Stick model as “BallStick_in1”, “BallStick_in2” and “BallStick_in3” for respectively one, two or three Stick compartments. This is a general naming scheme to denote models that can have one or more intra neuronal compartments relative to the other compartments. From the Ball&Stick model we derive the Fraction of Stick (FS) quantity, which is the sum of the volume fractions of the Stick compartments.

More recent, biologically inspired, models include Bingham-NODDI and CHARMED. The Bingham-NODDI model assumes that white matter consists of restricted intra-cellular and hindered extra-cellular water compartments, with the intra-cellular compartment capturing neurite orientation dispersion. From the Bingham-NODDI model we use the Fraction of Restricted (FR) quantity, the volume fraction of the restricted intra-cellular compartment. The CHARMED model assumes a tissue model of restricted intra-neuronal and hindered extra-neuronal water compartments, with the intra-neuronal compartment assuming a bundle of axons. Since CHARMED can be used with multiple intra-neuronal compartments we again denote these with the ‘ _in’ suffix. Here, we only use CHARMED with one intraneuronal compartment, denoted as “CHARMED_in1”. From the CHARMED model we use the Fraction of Restricted (FR) quantity, the volume fraction of the restricted intra-neuronal compartment. For implementation notes of these models see (Harms et al., 2017).

### 2.4 Software

All models and routines used in this study are implemented in a Python based GPU (graphical processing unit) accelerated toolbox, the Microstructure Diffusion Toolbox, MDT, which is freely available under an open source license at https://github.com/cbclab/MDT. We used the models and MCMC routine as implemented in MDT version 0.18.3. From this version onward, MDT automatically computes the FIM after every maximum like-lihood estimation operation and writes out the variances and covariances alongside the parameter estimates. Scripts for reproducing the results in this article can be found at https://github.com/robbert-harms/uncertainty_paper. All computations for this paper were performed on a single AMD Fury X graphics card.

### 2.5 Datasets

In this study we used simulated data and imaging data from two population studies. To illustrate the methods on a dataset with a clinically feasible, fast to acquire, acquisition scheme, we used data from the diffusion protocol pilot phase of the Rhineland Study (www.rheinland-studie.de). We refer to these datasets and acquisition schemes as *RLS-pilot*. To illustrate the methods on a dataset with a high-end, long acquisition time, acquisition scheme, we used data from the Human Connectome Project MGH-USC Young Adult study. We refer to these datasets and acquisition schemes as *HCP MGH*. For simulated data we used a single representative acquisition scheme from both the RLS-pilot and HCP MGH studies.

The RLS-pilot datasets were acquired on a Siemens Magnetom Prisma (Siemens, Erlangen, Germany) with the Center for Magnetic Resonance Research (CMRR) multi-band (MB) diffusion sequence (Moeller et al., 2010; Xu et al., 2013). These datasets had a resolution of 2.0 mm isotropic with diffusion parameters Δ = 45.8 ms, *δ* = 16.3 ms, TE = 90 ms and TR = 4500 ms, and with Partial Fourier = 6/8, MB factor 3, no in-plane acceleration with 3 shells of b = 1000, 2000, 3000 s/mm^2^, with respectively 30, 40 and 50 directions to which are added 14 interleaved b0 volumes leading to 134 volumes in total per subject. Additional b0 volumes were acquired with a reversed phase encoding direction which were used to correct susceptibility related distortion (in addition to bulk subject motion) with the topup and eddy tools in FSL version 5.0.9 (Andersson & Sotiropoulos, 2016). The total acquisition time is 10 min 21 sec. These three-shell datasets represent a relatively short time acquisition protocol that still allows many models to be fitted. From this dataset we used a single representative subject (v3a_1_data_ms20).

The HCP MGH datasets come from the freely available fully preprocessed dMRI data from the USC-Harvard consortium of the Human Connectome project. Data used in the preparation of this work were obtained from the MGH-USC Human Connectome Project (HCP) database (https://ida.loni.usc.edu/login.jsp). The data were acquired on a specialized Siemens Magnetom Connectom with 300 mT/m gradient set (Siemens, Erlangen, Germany). These datasets were acquired at a resolution of 1.5 mm isotropic with diffusion parameters Δ = 21.8 ms, *δ* = 12.9 ms, TE = 57 ms, TR = 8800 ms, Partial Fourier = 6/8, MB factor 1 (i.e. no simultaneous multi-slice), in-plane GRAPPA acceleration factor 3, with 4 shells of b = 1000, 3000, 5000, 10,000 s/mm^2^, with respectively 64, 64, 128, 393 directions to which are added 40 interleaved b0 volumes leading to 552 volumes in total per subject, with an acquisition time of 89 minutes. These four-shell, high number of directions, and very high maximum b-value datasets allow a wide range of models to be fitted. From these datasets we used a single representative subject (hcp_1003) in single subject illustrations and we used all 35 subjects in the group comparisons.

Since the CHARMED in1 model requires relatively high b-values (≥~6000 s/mm^2^), which are not present in the RLS-pilot datasets, we will only use the HCP MGH dataset when showing CHARMED_in1 results. Additionally, since the Tensor model is only valid for b-values up to about 1200 s/mm^2^, we only use the b-value 1000 s/mm^2^ shell and b0 volumes in maximum likelihood estimation and MCMC sampling of the Tensor model. All other models use all the data volumes.

For all datasets we created a white matter (WM) mask from the Tensor FA estimates and, using BET from FSL (Smith, 2002), a whole brain mask. The whole brain mask is used for MLE and MCMC sampling, whereas averages over the WM mask are used in model or data comparisons. For each dataset, voxel-wise SNR is estimated using only the unweighted (b0) volumes, by dividing the mean of the unweighted volumes by the standard deviation.

#### 2.5.1 Ground truth simulations

We additionally created simulated data to illustrate the effects of the signal-to-noise ratio (SNR) on the variance of the estimated parameters. We used a single representative acquisition scheme from both the RLS-pilot and HCP_MGH datasets (the acquisition schemes of subject v3a_1_data_ms20 and hcp 1003), and simulated data for each model. For each acquisition scheme and each model, we simulate 10000 voxels with random volume fractions in [0.2, 0.8], diffusivities in [5*e* − 11, 5*e* − 9] mm^2^/s, and orientations in [0, π]. From these, we created multiple copies with Rician noise (Gudbjartsson & Patz, 1995) of SNRs 5, 10, 20, 30, 40 and 50. We then fit and sample each model ten times to these simulated datasets and estimate the standard deviation using both the FIM and MCMC approach as described above. Per SNR we summarize the results of these ten trials as a mean standard deviation and its corresponding standard error of the mean.

#### 2.5.2 Group statistics

For the group statistics we computed Tensor FA and Bingham-NODDI FR and FR standard deviation maps on all 35 subjects using the MLE+FIM method. To be able to compare the subjects, we first registered the Tensor FA estimates to the FMRIB58_FA_1mm template using FLIRT and FNIRT from FSL (Andersson et al., 2010). Next, we used those registration templates to co-register the Bingham-NODDI FR and FR standard deviation maps.

With uncertainty maps available there are three methods to compute group statistics that are robust against subject-level artifacts. Method one, apply variance weighted averaging using the uncertainty estimates to down-weight voxels with a high standard deviation. This would automatically remove artifacts if these artifacts lead to high parameter uncertainties. Method two, exclude outlier subjects from the group statistic. Outlier subjects could be detected using the point estimates or using the uncertainty maps. Method three, use a combination of method one and two, i.e. computing weighted group estimates after removal of outliers.

To illustrate these three artifact reduction methods, we first computed a baseline statistic using a simple mean and standard deviation over all 35 subjects. We then used artifact reduction method one and used the FR standard deviation maps as weights in the variance weighted averaging. To apply artifact reduction method two and three, we created a new subgroup with only 30 subjects, where we manually removed five subjects (mgh_ - 1008, mgh_1009, mgh_1013, mgh_1017 and mgh 1032) that had a large white matter artifact over the corpus callosum. We then applied regular averaging and weighted averaging over these remaining 30 subjects.

As a comparison method between regular and weighted averaging we computed (*μ*_weighted_ − *μ*_regular_)/*μ*_regular_ and (*σ*_weighted_ − *σ*_regular_)/*σ*_regular_ as difference measure for the mean and standard deviation estimates between regular and weighted averaging.

## 3 Results

We begin by comparing the parameter estimates and parameter uncertainty estimates of MLE+FIM to the corresponding estimates from MCMC. Next, we investigate the effect of SNR on the parameter standard deviations using both simulated and imaging data. We end with a comparison of regular versus weighted averaging in group statistics.

### 3.1 Parameter distribution estimates

Figure 2 visually compares the results of MLE+FIM to those of MCMC, using the Bingham-NODDI Fraction of Restricted (FR) parameter, on a single subject from the RLS-pilot dataset. Comparing results of a single transverse slice shows high qualitative correspondence between the MLE and MCMC methods (figure 2A), with both the point estimates and corresponding standard deviations (stds.) in close resemblance. A single voxel illustration of the estimated Gaussian distributions (figure 2B) again shows a high degree of similarity, with both Gaussian fits capturing the characteristics of the MCMC sample distribution to a large degree.

**Figure 2:**
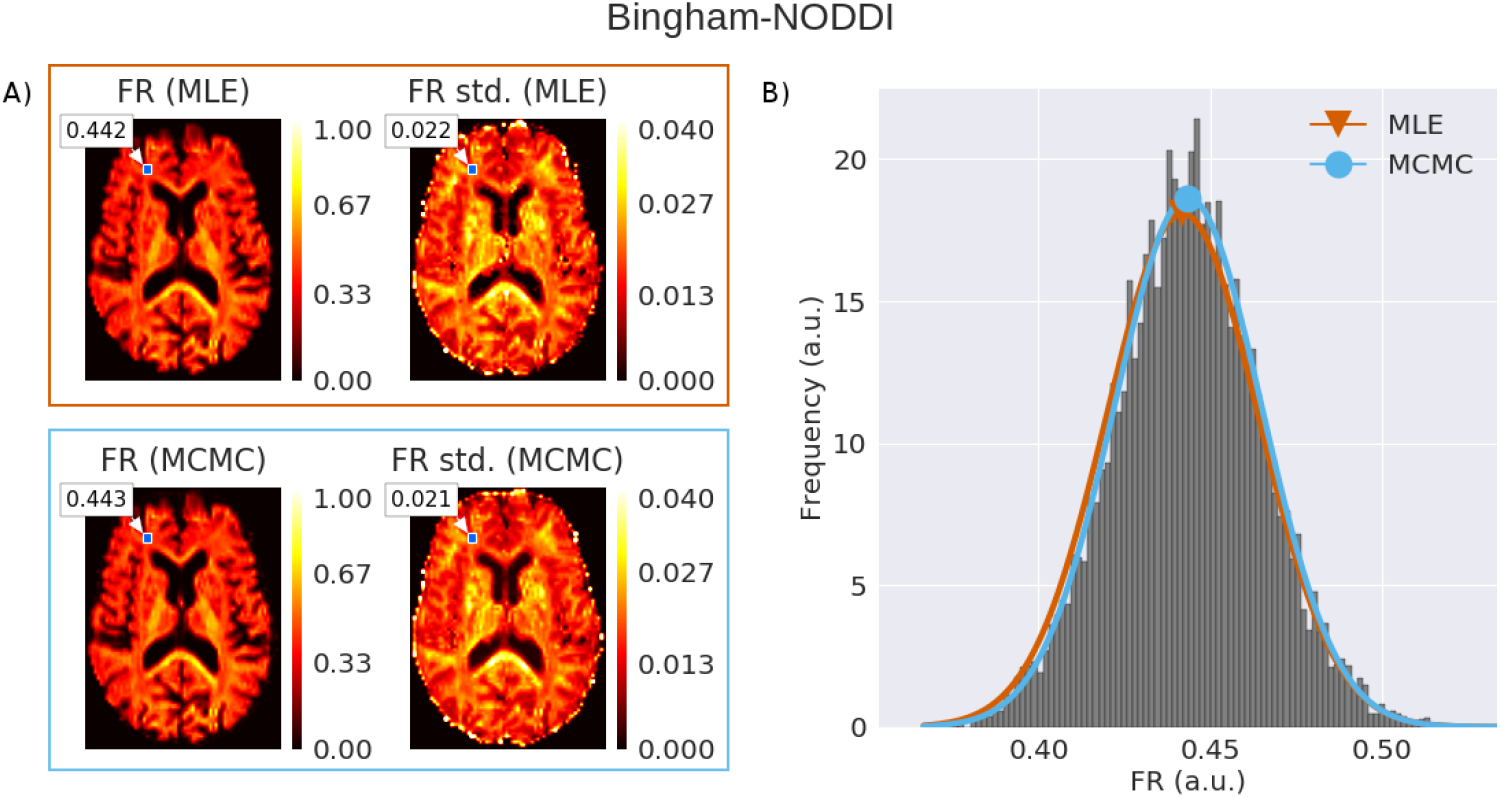
A) Visual comparison of parameter and standard deviation uncertainty maps between the Maximum Likelihood Estimation (MLE) and Markov Chain Monte Carlo (MCMC) methodologies for the Bingham-NODDI Fraction of Restricted (FR) on an RLS-pilot dataset. B) Histogram of the 20 thousand MCMC samples of the highlighted voxel in figure A, with in red and blue the fitted Gaussian distributions of, respectively, the MLE and MCMC methodologies.

To further quantify the correspondence between the MLE and MCMC methodologies, we created scatter plots between the MLE and MCMC estimates of both the point estimate and standard deviation estimate. This was performed over a white matter mask for a single subject from both the HCP MGH and RLS-pilot datasets. Figure 3 shows Bingham-NODDI FR mean and standard deviation scatter plots. The FR point estimates are very tightly confined to the identity line, illustrating a high degree of correspondence in the point estimates from MCMC and MLE. The variation of point estimates along the diagonal corresponds to variation of FR values over the white matter mask, ranging between roughly 0.3 and 0.7. The std. estimates between the MLE and MCMC methodologies again show a high correspondence, although the off-diagonal spread in the std. plot is visibly larger than that in the point estimate plot. There is also some clipping visible in the std. plot, with MLE estimating a zero std. while MCMC provides a range of values. This is mostly due to very low point estimates (near zero), at which point the FIM is no longer applicable. The blue-green-yellow-red coded points in both plots account for 97-99.5% of the voxels and the purple points account for the remaining fraction of outliers. The std. estimates for the HCP MGH data are clearly lower than for the RLS-pilot data, confirming an expected higher precision (lower uncertainty) of point estimates based on more dMRI data-points.

**Figure 3:**
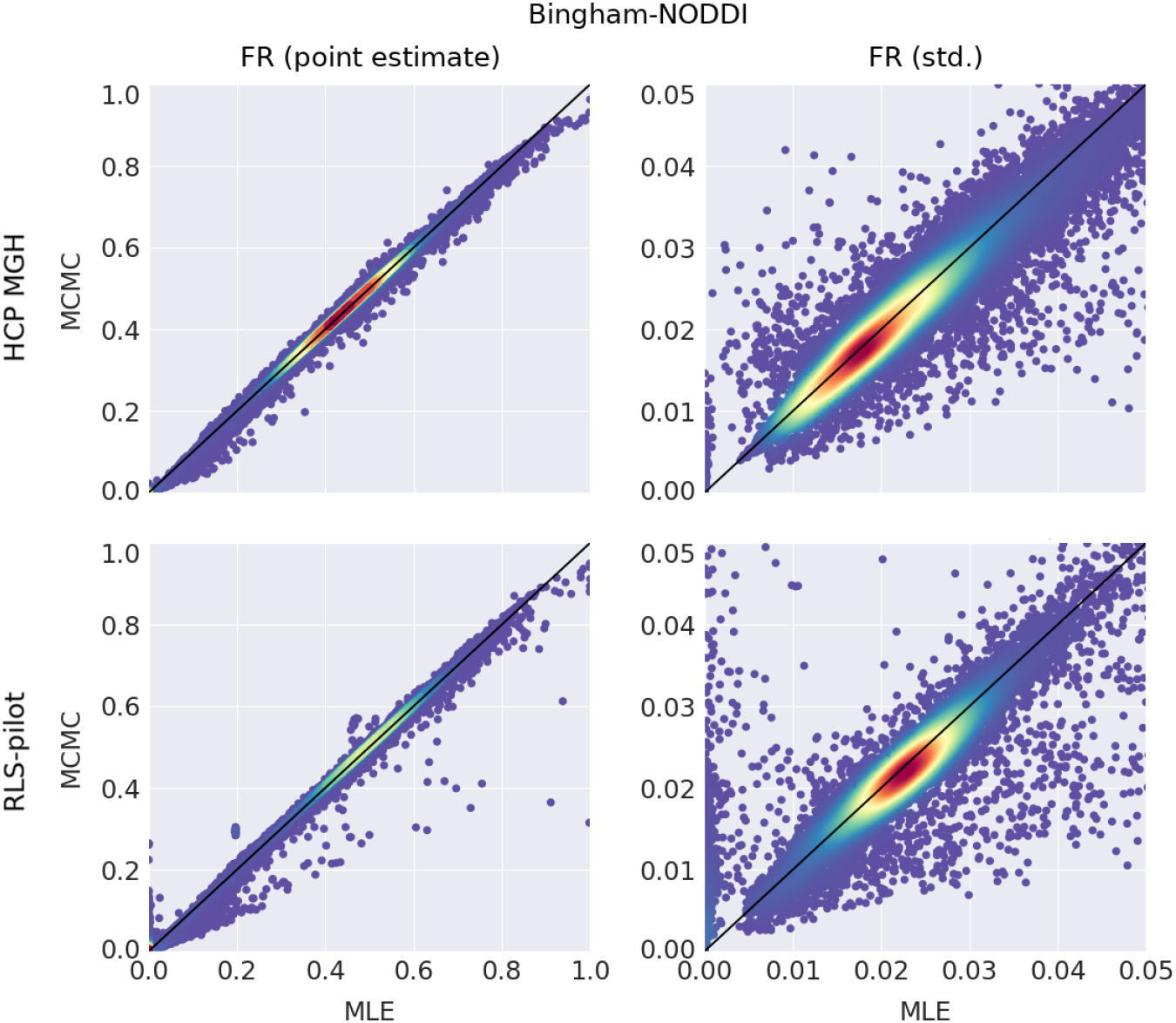
Scatter plots comparing Maximum Likelihood Estimation (MLE) and Markov Chain Monte Carlo (MCMC) point estimates (left column) and standard deviations (right column) for the Bingham-NODDI Fraction of Restricted (FR) values over a white matter mask for both a complex, long acquisition time HCP MGH dataset and a clinically feasible RLS-pilot dataset. Plots are color coded using a kernel density estimate (a.u) from purple (low density) to red (high density). Purple points correspond to a small percentage (0.5-3%) of the data (c.f. Table 1).

To investigate the correspondence in MCMC and MLE uncertainty estimates for a larger number of models, figure 4 shows scatter plots for multiple microstructure models. Parameter point estimate comparisons are not shown here, but are generally in correspondence to a very high degree. Across all models and data, except for the CHARMED in1 model fit on RLS-pilot data, MCMC and MLE uncertainty estimates are in high correspondence and located close to the identity diagonal. A relatively large off-diagonal variance in standard deviation estimates is visible in the CHARMED_in1 FR parameter on the RLS-pilot data. This is expected as the RLS-pilot dataset is not well suited for the CHARMED in1 model due to too low b-values (the CHARMED_in1 model requires b-values ≤ 6000s/mm^2^). Standard deviation estimates for CHARMED in1 on the HCP MGH data are not only much more tightly confined to the identity diagonal, the std. estimates themselves are also about a factor two lower. A large spread to the right is also visible in the Ball&Stick_in3 results. This might be related to MLE choosing a different mode and is perhaps solved using model selection. There is also again some clipping visible, with MLE providing a zero std. with voxels with a very low point estimate.

**Figure 4:**
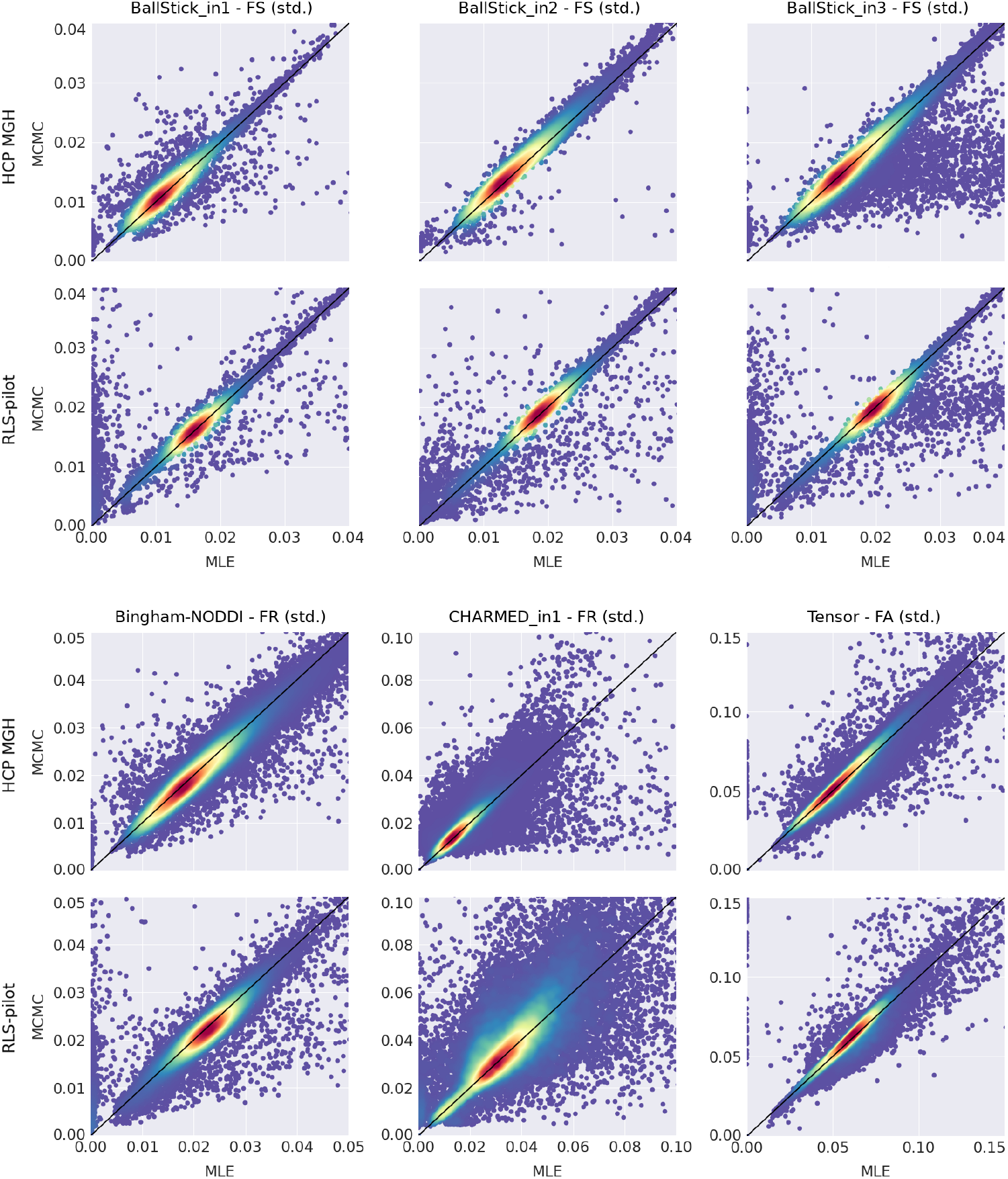
Scatter plots comparing Maximum Likelihood Estimation (MLE) and Markov Chain Monte Carlo (MCMC) standard deviations for multiple models over a white matter mask for both an HCP MGH and an RLS-pilot dataset. Acronyms are Fraction of Stick (FS), Fraction of Restricted (FR) and Fractional Anisotropy (FA). Plots are color coded using a kernel density estimate (a.u) from purple (low density) to red (high density). Purple points correspond to a small percentage (0.5-3%) of the data (c.f. Table 1).

Irrespective of the method (MCMC or MLE+FIM), the std. estimates on the RLS-pilot data are always higher than the corresponding estimates on the HCP MGH data, once again confirming the expected higher precision on datasets with a larger number of direction. Conversely, one would expect higher complexity models (i.e. models with more compartments and more parameters to fit) to have higher uncertainty when fitted on the same data. This is indeed illustrated by the Ball&Stick in{1,2,3} results, were we see an increasing estimated standard deviation for an increasing number of Sticks, within each of the HCP MGH and RLS-pilot datasets. Finally, Tensor FA standard deviations are about a factor two higher than those of the other models. This is probably related to Tensor FA being a compound parameter.

To quantify correspondence in the MCMC and MLE std. estimates in the scatter plots, table 1 shows the percentage of voxels for which the difference between the MLE and MCMC variances is less than two standard deviations from the mean difference. We note an average similarity of ~98.7% across six models and two datasets, even including the 97.9% similarity for the CHARMED in1 model fit on RLS-pilot data. Table 2 compares runtimes between the MLE with the FIM and the MCMC methodologies, measuring the time between loading the data and writing the results. Averaged over six models and two subjects, the GPU-optimized MLE and FIM together compute approximately 38 times faster than GPU-optimized MCMC.

**Table 1:** For each model and dataset the percentage of voxels where the difference between the parameter stds. from the FIM and of MCMC are within two standard deviations from the mean difference. These percentages correspond to the red/yellow high densities in figure 4.

|  | **HCP MGH** | **RLS-pilot** |
| --- | --- | --- |
| Ball&Stick_in1 | 99.5% | 98.8% |
| Ball&Stick_in2 | 99.9% | 99.4% |
| Ball&Stick_in3 | 98.8% | 97.5% |
| Bingham-NODDI | 98.9% | 98.7% |
| CHARMED_in1 | 98.6% | 96.9% |
| Tensor | 99.0% | 97.9% |

**Table 2:** Runtime comparison between the two methodologies for computing parameter statistics, Maximum Likelihood Estimation (MLE) with the Fisher Information Matrix (FIM) and Markov Chain Monte Carlo (MCMC) sampling, for six different models and using a single representative subject from both the HCP MGH and the RLS-pilot datasets. Reported run times are over the entire brain mask and are in units of (h:m:s), with next to it the relative speed advantage of the MLE + FIM over MCMC.

|  | **HCP MGH** |  |  | **RLS-pilot** |  |  |
| --- | --- | --- | --- | --- | --- | --- |
|  | *MLE + FIM* | *MCMC* | rel. | *MLE + FIM* | *MCMC* | rel. |
| Ball&Stick_in1 | 00:01:49 | 01:21:55 | 45x | 00:00:30 | 00:20:49 | 42x |
| Ball&Stick_in2 | 00:04:32 | 02:33:18 | 34x | 00:01:08 | 00:42:36 | 38x |
| Ball&Stick_in3 | 00:13:01 | 07:00:51 | 32x | 00:03:19 | 01:53:33 | 34x |
| Bingham-NODDI | 02:06:19 | 111:32:52 | 53x | 00:28:19 | 26:11:47 | 56x |
| CHARMED_in1 | 02:09:49 | 53:34:47 | 25x | 00:21:53 | 07:49:55 | 21x |
| Tensor | 00:02:41 | 01:59:07 | 44x | 00:02:18 | 01:02:11 | 27x |

### 3.2 Effect of SNR on parameter variances

Lower SNR per data point (i.e. single diffusion volume) is expected to lead to higher uncertainty in fitted parameter estimates. This issue is of extra importance in brain dMRI by the fact that SNR is non-uniform over the brain, especially in modern high number-of-channel phased array RF-coils. In order to assess the effect of SNR on parameter variances, figure 5 compares an estimate of SNR, its reciprocal, and the parameter standard deviation estimates of multiple white matter models on a single HCP MGH dataset. We observe a decreased SNR in the center of the brain and an increase of SNR towards the periphery. A very similar gradient can be observed in the standard deviation maps, with a decrease in parameter standard deviations towards the periphery. As in the previous results, we observe an increase in standard deviations for an increased number of Sticks in the Ball&Stick_ -in{1,2,3} models, and Tensor FA standard deviations are about a factor two higher than the other standard deviation estimates.

**Figure 5:**
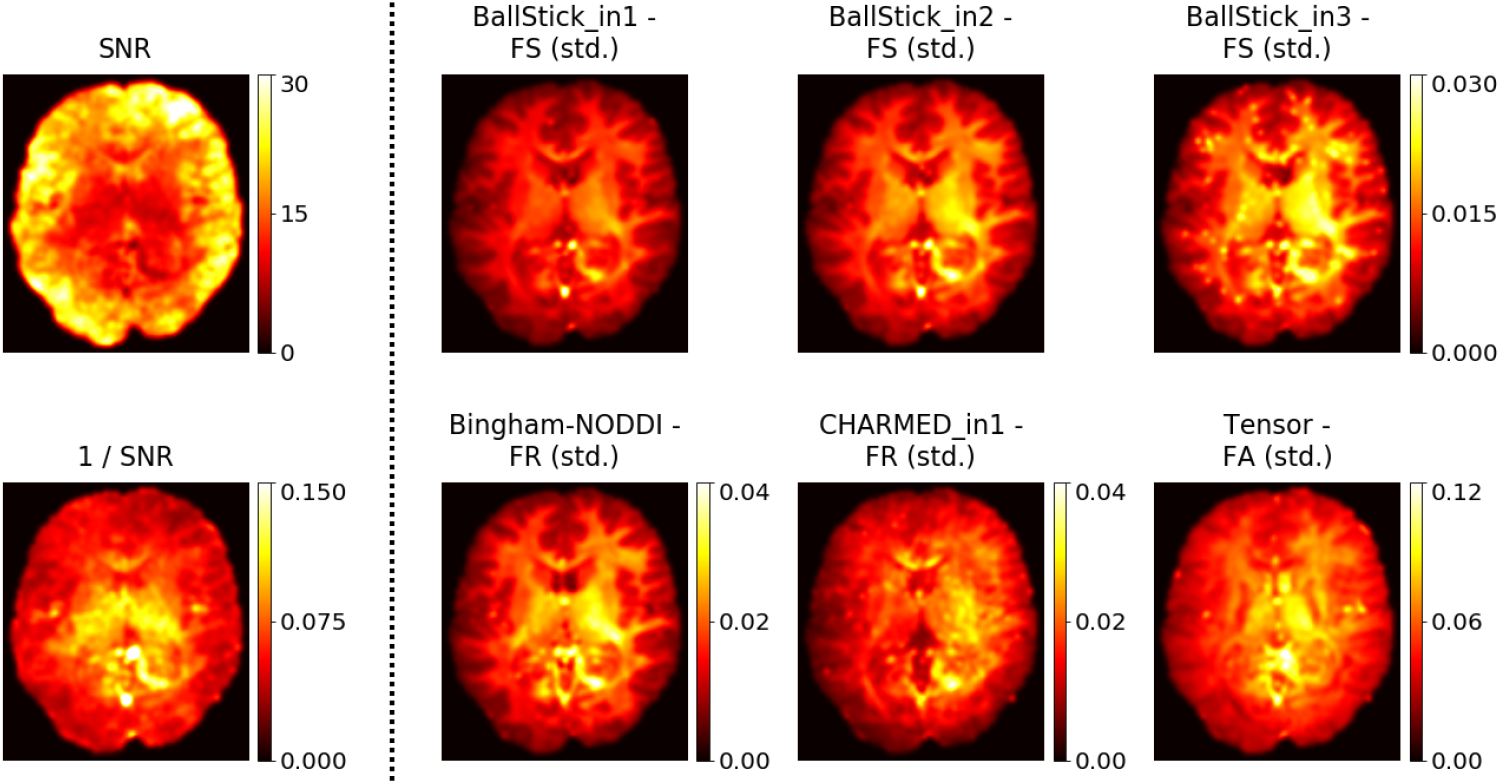
Illustration of the effect of Signal to Noise ratio (SNR) on parameter standard deviation estimates (using the MLE methodology), for a single HCP MGH subject (subject 1003). Maps are slightly smoothed with a 3d Gaussian filter (σ = 1*voxel*). Parameter acronyms are Fraction of Stick (FS), Fraction of Restricted (FR) and Fractional Anisotropy (FA).

To further compare SNR and standard deviation estimates, figure 6 plots SNR versus parameter standard deviations, for both simulated data and imaging data. In general, we observe an inverse relationship between SNR and standard deviation, where an increase in SNR leads to an decrease in parameter std. estimates. Standard deviations on RLS-pilot data are always higher than corresponding estimates on HCP MGH data, except for the imaged data analysis at an SNR of 5, where the RLS-pilot dataset has a lower standard deviation. For lower SNR (< 10), MLE std. estimates are slightly higher than the MCMC estimates. For higher SNR (> 10), the MLE and MCMC standard deviation estimates quickly converge, except for Ball&Stick_in2, Ball&Stick_in3 and Tensor estimates on the RLS-pilot dataset, where MLE standard deviations stay higher than those from MCMC. For the HCP MGH dataset, results are consistent between simulated and imaging data, with differences within the Standard Error of the Mean (SEM). Results on the RLS-pilot dataset are generally also consistent, except for an SNR of 5, where imaging data results are lower than those on simulated data. We finally observe that the standard error of the mean is generally higher for the simulated data compared to the imaging data, especially for lower SNR.

**Figure 6:**
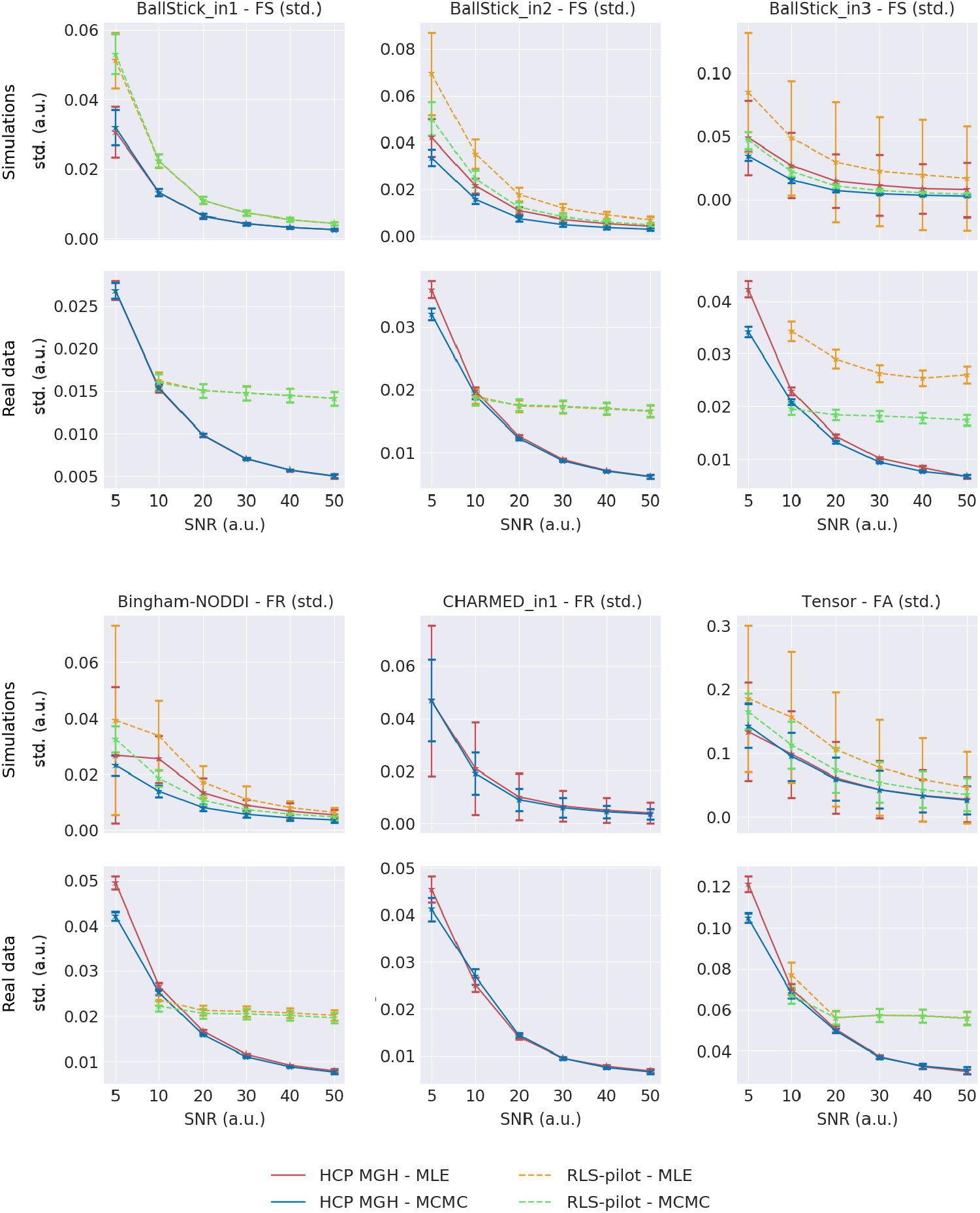
Effect of SNR on parameter standard deviations for simulated data and imaging data. Simulation results are over 10000 simulated voxels per SNR with a standard error of the mean (SEM) as error bar over 10 optimization and sampling trials. Real data results are for 10 subjects of the HCP MGH and 10 subjects of the RLS-pilot datasets, with SNR estimated as mean(b0_volumes)/std(b0_volumes).

### 3.3 Group statistics

Figure 7 shows Bingham-NODDI FR results of three subjects of the HCP MGH dataset after co-registration, to illustrate the behavior of standard deviations in regions of white matter acquisition artifacts. The first subject (top row) has a clear artifact across the genu of the corpus callosum, perhaps due to incomplete fat saturation. This artifact is visible in both the mean parameter estimates and the standard deviation estimates. The second subject (middle row) shows a patch of relatively large standard deviations in and near the splenium of the corpus callosum, without an easily detectable alteration in the mean parameter map. For comparison, we show a third subject (bottom row) at the same contrast scaling, with no visible artifacts or alterations in either the mean or standard deviation estimates. This figure illustrates that parameter std. maps can play a role in detecting biased estimates resulting from imaging artifacts. In particular, artifacts which may not always be detectable in the parameter maps themselves.

**Figure 7:**
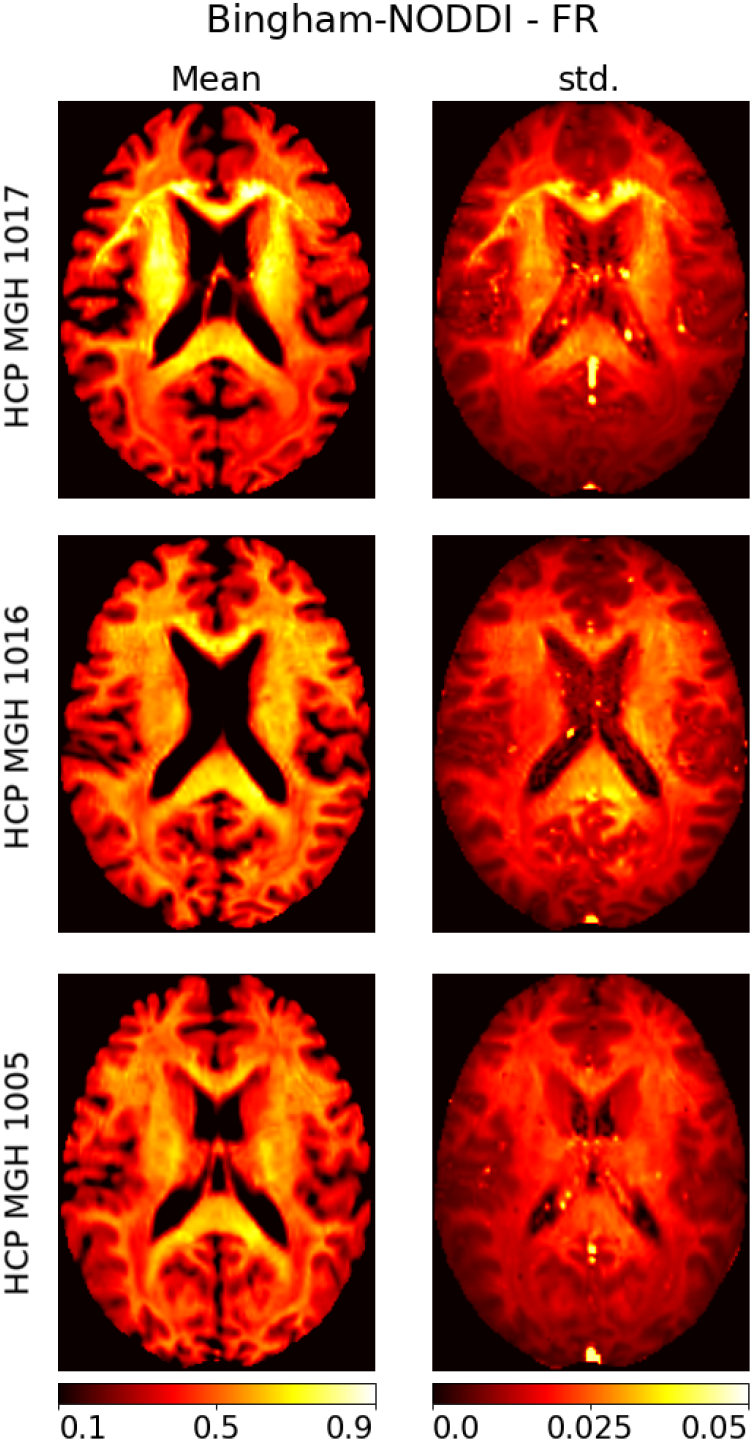
Illustration of artifacts in the HCP MGH datasets using the Bingham-NODDI Fraction of Restricted (FR) mean and standard deviation (std.) estimates from the MLE methodology. In the top row, estimates for HCP MGH subject 1017, with an artifact across the corpus callosum. In the middle row, estimates for HCP MGH subject 1016 with increased standard deviation estimates near a ventricle. In the bottom row, estimates for HCP MGH subject 1016 with no artifacts visible in the mean or standard deviation map.

Figure 8 shows four group statistic estimates, a regular (baseline) and three statistics using the three mentioned artifact reduction methods using the parameter variances. To reiterate, these were method one, a weighted average on all 35 subjects, method two, remove outlier subjects and apply regular averaging and method three, a weighted average with outlier subjects removed. Between regular and weighted averaging we computed a percentile difference map over a white matter mask to highlight the differences in estimates of both the group mean and group standard deviations.

**Figure 8:**
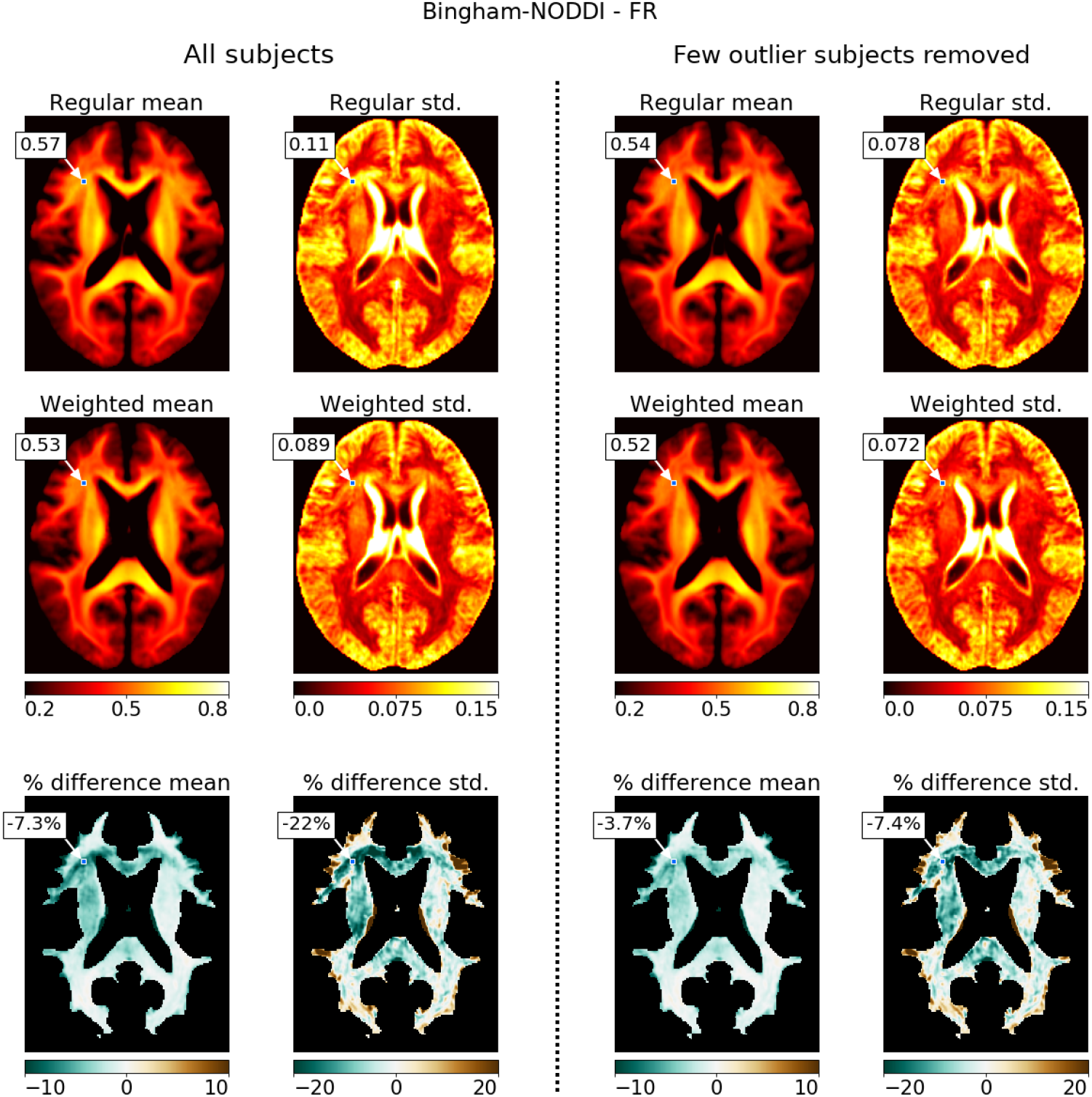
Group averages of Bingham-NODDI Fraction of Restricted (FR) estimates using the HCP MGH data, once over all 35 subjects (left two columns) and once over only 30 subjects where 5 outlier subjects have been removed (right two columns). First row, the regular mean and standard deviation, second row, the variance weighted mean and standard deviations, final row, percentage difference between regular and weighted averages. Point estimates and variances were computed using the MLE methodology.

For both the all-subjects and outliers-removed subject groups, the variance weighted mean is approximately lower across the artifact above the corpus callosum and, to a lesser degree, over the left internal and external capsules. For both groups, standard deviation estimates vary more between regular and weighted averaging, with a lower weighted average across the white matter artifact, equal values in most of the white matter and higher estimates near the border with gray matter. Group statistics with a few out-lier subjects removed give lower averages and lower standard deviations for both weighted and regular averaging. Removing the outlier subjects brings the regular and weighted averages closer to each other, with percentile differences dropping by at least half.

The white matter artifact is most present in the regular average over all subjects (baseline), followed by regular averaging over the reduced group (artifact reduction method two), then by weighted averaging over all subjects (artifact reduction method one), and the artifact is least present in weighted averaging over the reduced group (artifact reduction method three).

## 4 Discussion

We evaluated parameter variance estimates as a quantification of parameter uncertainties. We compared standard deviation estimates from Maximum Likelihood Estimation (MLE) plus the Fisher Information Matrix (FIM) to those of Markov Chain Monte Carlo (MCMC) sampling and showed that both results are identical in ~98.7% of the voxels. In terms of computer processing time, the estimates of MLE+FIM computed about 38x faster than those of MCMC. We then showed how data complexity and the signal-to-noise ratio can affect the parameter variances. Finally, we illustrated how the parameter variances can be applied in group studies to identify and downweight the effect of outliers, thereby decreasing the variance in group estimates, leading to an increase in statistical power of group studies.

### 4.1 Comparison of the FIM and MCMC

In general, we noted a close correspondence between the parameter distribution estimates from the FIM and those from MCMC sampling, with an average similarity of ~98.7% across six models and two datasets. Compared on runtime, computing MLE+FIM is about 38x faster than the use of MCMC, for comparable results.

We made the explicit assumption that the parameter posterior distributions would follow a Gaussian distribution with a single mode. Theoretically, only a symmetrical distribution with a single mode would have an equal mode and mean. Therefore, if the MLE point estimate, which attempts to find the mode of the posterior, and the MCMC point estimate, which was computed here as the mean of the sample distribution are equal, then this is evidence towards symmetric single mode posteriors. Since our results from the FIM and MCMC were highly comparable (i.e. up to 98.7% of points estimates were indeed nearly equal), the Gaussian assumption is often confirmed. In the remaining 1.3% of the voxels, it could either be that the parameter posteriors were not fully Gaussian distributed, or that the posterior distributions were multi-modal. In the case of a multi-modal distribution, the FIM will give variance estimates around a single mode only, the mode found by the maximum likelihood routine. Our current MCMC methodology would provide an average and variance over all modes. To proper deal with multi-modal distributions when using MCMC, would require fitting a multi-modal normal distribution to the MCMC samples. If the parameters are not normally distributed, like for example near parameter boundaries or with skewed posterior’s, the FIM no longer applies and MCMC would require different post-processing of the samples.

Compared on signal-to-noise (SNR), we note that the FIM provides higher standard deviation estimates at low SNR (< 10) when compared to MCMC. These differences are small and quickly vanish for SNR ≥ 10. This follows results from astrophysics, where they recommend a minimum SNR of 10 to compute variances using the FIM, in gravitational wave assessments (Rodriguez et al., 2013).

In general, both the FIM and MCMC give comparable answers and both can be used for computing parameter standard deviations estimates to compute uncertainty. The only essential difference is one of computation time, computing a maximum likelihood point estimate together with the FIM is about 38x faster than using MCMC. This was expected, MCMC is generally known to be a time-consuming process, even when run on a GPU (Harms & Roebroeck, 2018). MLE on the other hand can be applied very efficiently using a GPU (Harms et al., 2017) and computing the FIM requires only a few extra function evaluations (dependent on the number of parameters, see Appendix A).

### 4.2 Effects on estimates of the standard deviations

There are several model and data characteristics that can affect standard deviation estimates, like data complexity, derived parameter maps and the signal-to-noise ratio. In general, these effects apply equally to both the FIM and MCMC.

Concerning data dependency, as expected, standard deviation estimates on the RLS-pilot dataset are generally higher than those on the HCP MGH dataset, reflecting a decrease in point estimate uncertainties with more data points. The same holds for the relatively large standard deviations in the Tensor Fractional Anisotropy (FA) estimates, since for the Tensor model we used only the data volumes with a low b-value.

A higher variance can additionally be observed for parameter maps which are not estimated directly but derived from the estimated parameters. This makes the variance of such derived parameters maps also a function of multiple variances, often leading to a higher total variance. This can for example be observed in the Tensor FA measure. The same compound effect could apply to the variance of the Fraction of Stick (FS) of the Ball&Stick models. For an increasing number of Sticks, the variance in FS is also a function of multiple volume fractions, which could increase the total variance.

For all models, parameter standard deviations are influenced by the signal-to-noise (SNR) ratio of the data, with a low SNR (< 10) leading to a large increase in standard deviations. Both shown in real and simulated data, the effect of SNR on the standard deviation estimates seems to be more gradual after an SNR ≥ 20.

### 4.3 Artifact detection

The computed parameter standard deviations (either from the FIM or MCMC) could be used as a tool for detecting acquisition artifacts. In one provided example (figure 7 top row), an artifact in the white matter was visible in both the parameter estimate and in the standard deviation as a patch of high intensity voxels. In another example (figure 7 middle row), a patch of high intensity voxels was visible in the standard deviation estimate but not in the parameter estimate itself. As such, standard deviation maps have the potential to be more sensitive in detecting white matter artifacts than point estimate maps themselves. A promising future development could be to include these standard deviation maps into quality control frame-works (Bastiani et al., 2019; Liu et al., 2010; Oguz et al., 2014).

### 4.4 Increasing power in group studies

By weighing down voxels with a high standard deviation, weighted averaging can reduce the effect of white matter artifacts, approach lower and more accurate estimates of group variances and increase power of group statistics. In theory, if the within group datapoints are distributed with the same mean, variance weighted averaging promises the lowest possible variance in the group mean. We observe this in large parts of the white matter where weighted averaging lowers the variance in the group average as expected, thereby indirectly increasing power in group comparisons.

We have shown that some white matter artifacts are visible in the parameter standard deviation maps as patches of relatively large standard deviations. Since variance weighted averaging automatically reduces the effects of outliers whenever they have a large variance, variance weighted averaging automatically reduces the presence of artifacts. Even after removing a few subjects with a similar artifact, white matter averaging still reduces the presence of what appears to be a lower-expressed artifact in the remaining subjects. Due to this mechanism, subjects no longer need to be excluded from analysis, thereby improving the power of one’s study.

Near the gray-white matter border we noticed some voxels where weighted averaging provides a higher variance than regular averaging. Theoretically, weighted averaging only predicts lower standard deviations if the points are distributed with the same mean. Misalignment between subjects can cause a single voxel to contain white matter for one subject and gray matter for another subject. Parameter estimates on such voxels will then be distributed with a different mean, leading to a higher group standard deviation when applying weighted averaging. This could be considered to be desirable, since such misalignment should not lead to high certainty group results and is therefore downweighted by the weighted averaging. In other words, the weighted group standard deviation could diagnose alignment errors in group studies.

We note that although weighted averaging is shown here over subjects, weighted averaging can also be applied within subjects. For example, when averaging voxels over a white matter tract. In essence, weighted averaging can be applied in all cases where variances of an estimate are available. In the future this could be applied to tract based microstructure or tractometry studies (Bells et al., 2011), for tract based summary statistics with a lower variance.

## 5 Conclusions and recommendations

Considering the advantages in processing time and close correspondence to Markov Chain Monte Carlo estimates, we recommend the use of the Fisher Information Matrix theory to quantify the uncertainties in parameter estimates. In individual subjects, the parameter standard deviations can help in detecting white matter artifacts as patches of relatively large standard deviations. In group statistics, we recommend using the parameter standard deviations by means of variance weighted averaging. Doing so can reduce the overall variance in group statistics and reduce the effect of data artifacts without discarding data from the analysis. Both these effects can lead to a higher statistical power in group studies.

## 6 Acknowledgements

RLH, FJF, SS and AR were supported by an ERC Starting Grant (MULTI-CONNECT, #639938), AR was additionally supported by a Dutch science foundation (NWO) VIDI Grant (#14637). This paper reflects only the author’s views and the European Union is not liable for any use that may be made of the information contained therein. Data collection and sharing for this project was provided, in part, by the MGH-USC Human Connectome Project (HCP; Principal Investigators: Bruce Rosen, M.D., Ph.D., Arthur W. Toga, Ph.D., Van J. Weeden, MD). HCP funding was provided by the National Institute of Dental and Craniofacial Research (NIDCR), the National Institute of Mental Health (NIMH), and the National Institute of Neurological Disorders and Stroke (NINDS). HCP data are disseminated by the Laboratory of Neuro Imaging at the University of California, Los Angeles. Collectively, the HCP is the result of efforts of co-investigators from the University of California, Los Angeles, Martinos Center for Biomedical Imaging at Massachusetts General Hospital (MGH), Washington University, and the University of Minnesota. Data collection and sharing for this project was provided, in part, by the Rhineland Study (Principal Investigator: Monique M.B. Breteler, M.D., Ph.D.; German Center for Neurodegenerative Diseases (DZNE), Bonn).

## Appendix A Numerical Hessian

To compute the Hessian we use a numerical differentiation routine with multiple step sizes and extrapolations to provide an estimate with a 𝒪(*h*^6^) order of accuracy. For a single step size vector **d**, we compute each element of the Hessian using a second order Taylor expansion central difference,

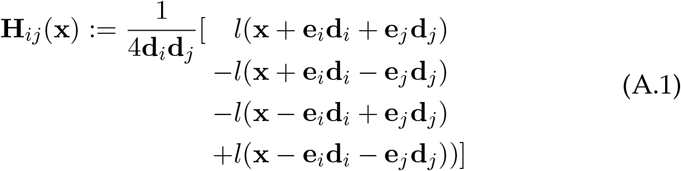

where 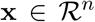 is the parameter vector, *l*(x) is the log-likelihood function and e_*k*_ is a zeros vector with only element *k* set to one. We evaluate the Hessian multiple times with exponentially diminishing steps and with the largest step size chosen such that x ± d is within bounds and d is within predefined upper and lower limits. In this work we evaluate the Hessian for five different step sizes d with each step half the previous step. We then apply Richardson extrapolation (Burg & Erwin, 2009) twice to produce three estimates with a sixth order of accuracy. These three approximations we extrapolate again using Wynn’s epsilon algorithm (Weniger, 1991) to arrive at a single final estimate.

## Appendix B Uncertainty propagation

This appendix provides two illustrations of uncertainty propagation, one example using Ball&Stick Fraction of Stick and one example using Tensor Fractional Anisotropy.

Uncertainty propagation of the Ball&Stick Fraction of Stick can be defined as follows. For a two Stick Ball&Stick model, the Fraction of Stick is defined as:

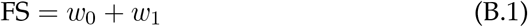

The analytical gradient of this function is given by:

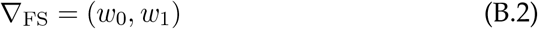

The covariance matrix of the weights can be defined as:

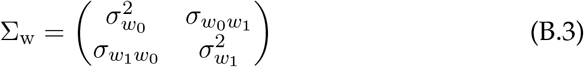

with 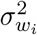 denoting the variance of weight *w*_*i*_, and 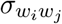 denoting the co-variances of weights *w*_*i*_ and *w*_*j*_. When evaluated, these quantities are taken from the covariance matrix provided by the FIM.

Using equation 6, we can write the uncertainty propagation as:

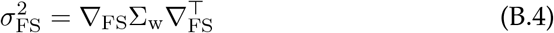

which simplifies to:

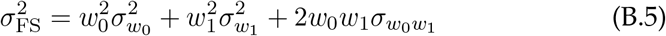

By evaluating expression B.5 using the point estimates, variance estimates and covariance estimates of the weights, we can compute the variance in the FS metric.

Uncertainty propagation of Tensor FA is slightly more complex considering FA is not a linear function of its inputs. The Tensor FA can be defined in terms of the three Tensor diffusivities (the eigenvalues of the diffusion Tensor) as:

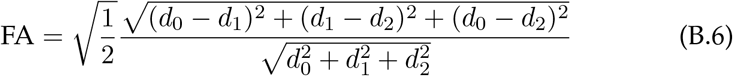

The derivative of FA with respect to the first diffusivity can be written as:

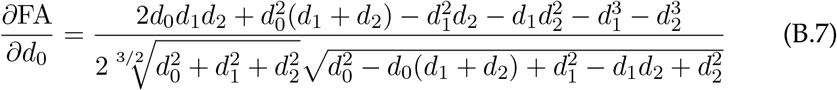

and similar derivatives can be derived for the second and third diffusivity by suitable permutations of the diffusivity indices. The analytical gradient of FA, ∇_FA_ can now be defined as:

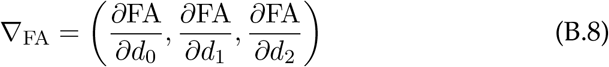

The covariance matrix of the diffusivities can be defined as:

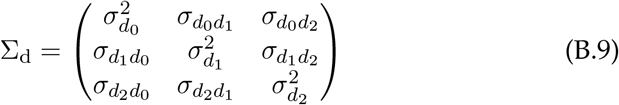

with 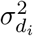 denoting the variance of diffusivity *d*_*i*_, and 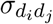 denoting the co-variances of diffusivities *d*_*i*_ and *d*_*j*_.

Using equation 6, we can define the uncertainty propagation of FA as:

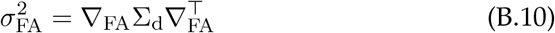

By evaluating expression B.10 using the point estimates of the diffusivities together with the corresponding variance and covariance estimates from the FIM, we can compute the propagated variance in the FA metric.

