## Supplementary data for "Fast quantification of uncertainty in non-linear diffusion MRI models for artifact detection and more power in group studies"

to the article

### 1 Parameter distribution estimates

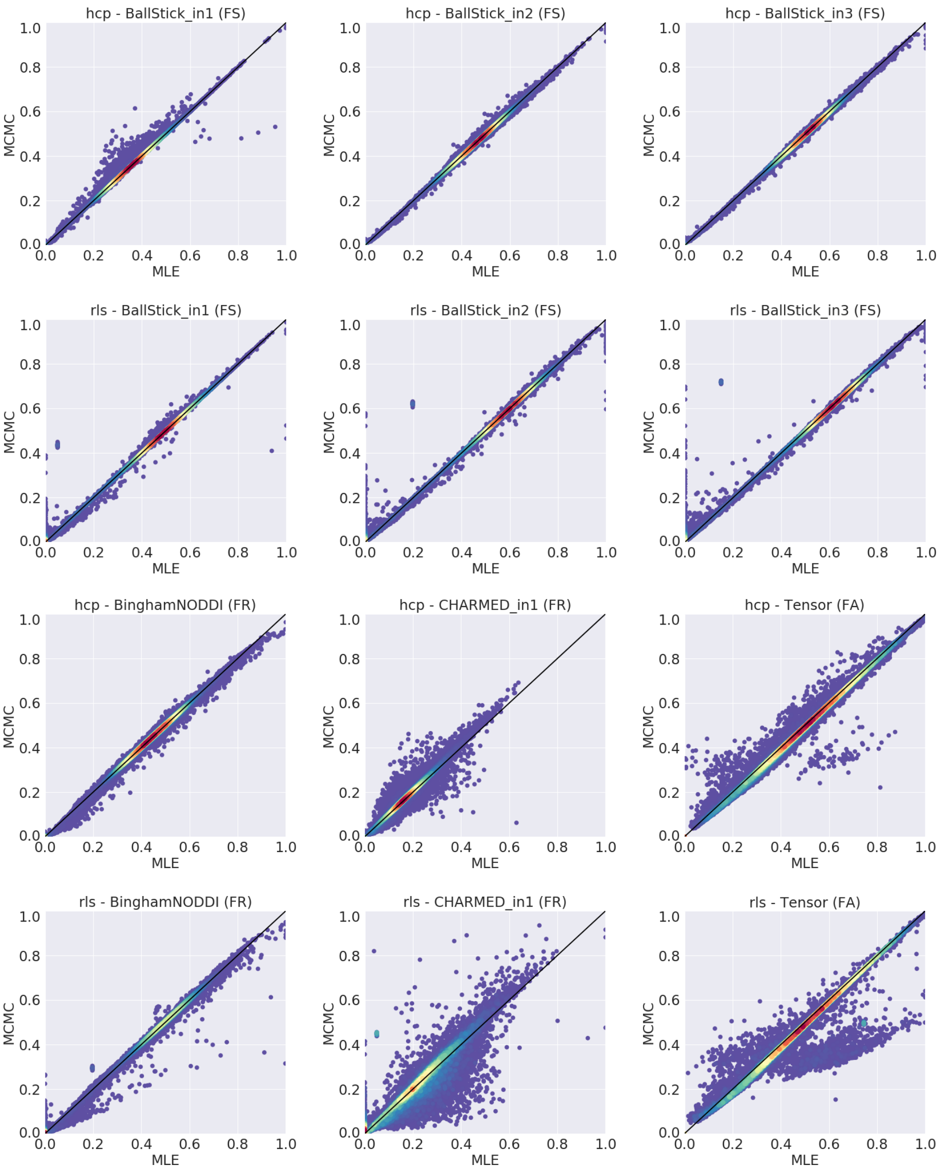


Figure 1: Scatter plots comparing Maximum Likelihood Estimation (MLE) and Markov Chain Monte Carlo (MCMC) parameter point estimates for multiple models over a white matter mask for both an HCP MGH and an RLS-pilot dataset. Acronyms are Fraction of Stick (FS), Fraction of Restricted (FR) and Fractional Anisotropy (FA). Plots are color coded by density (a.u) from blue (low density) to red (high density). The blue outlier dots generally constitute ¡ 2% of the data.
